# Bridging Domains in Chronic Lower Back Pain: Large Language Models and Ontology-driven Strategies for Knowledge Graph Construction

**DOI:** 10.1101/2024.03.11.584505

**Authors:** Paul Anderson, Damon Lin, Jean Davidson, Theresa Migler, Iris Ho, Cooper Koenig, Madeline Bittner, Samuel Kaplan, Mayumi Paraiso, Nasreen Buhn, Emily Stokes, Tony Hunt, Glen Ropella, Jeffrey Lotz

## Abstract

Link prediction and entity resolution play pivotal roles in uncovering hidden relationships within networks and ensuring data quality in the era of heterogeneous data integration. This paper explores the utilization of large language models to enhance link prediction, particularly through knowledge graphs derived from transdisciplinary literature. Investigating zero-shot entity resolution techniques, we examine the impact of ontology-based and large language model approaches on the stability of link prediction results. Through a case study focusing on chronic lower back pain research, we analyze workflow decisions and their influence on prediction outcomes. Our research underscores the importance of robust methodologies in improving predictive accuracy and data integration across diverse domains.

## 1 Introduction

To understand the complex interactions within biological systems, dynamic and comprehensive approaches must be used to investigate relationships between diverse entities such as genes, diseases, and even between disparate fields. Relationship discovery and link prediction have emerged as crucial components advancing our knowledge of health, disease mechanisms, and complex systems[9]. Link prediction algorithms can discover potential associations between biological entities, allowing researchers and clinicians to identify novel connections that may have been previously unnoticed[1]. Chronic pain is a complicated and pervasive health challenge that moves beyond traditional disciplinary boundaries, requiring a transdisciplinary approach for comprehensive understanding and effective management. This complex condition involves interactions among biological, psychological, social, and environmental factors, posing significant challenges to researchers and healthcare professionals, who are often siloed and isolated from other fields[19,7].

The broad context of chronic pain requires the integration of insights from diverse fields such as neuroscience, psychology, sociology, and public health, while those fields are commonly disincentivized from seeking interdisciplinary connections. Link prediction, with its ability to unveil relationships within complex systems, holds immense potential in addressing this current research environment. It could aid in identifying novel connections between genetic predispositions and environmental factors, correlations of psychological and social variables, and predicting the impact of interventions on individuals with chronic pain. By leveraging link prediction, researchers could discover novel insights that may allow for more targeted and personalized approaches to pain management. We propose that the use of Large Language Models (LLMs) to process literature from specific domains into knowledge graphs will allow for the automation of interdisciplinary discovery and novel connections, the key to uncovering insights in this challenging condition.

Link prediction is specifically focused on predicting future or missing connections in a network. It aims to uncover potential relationships between nodes that are not explicitly stated in the existing network. Previous areas where this type of novel connection finding has been useful include public health, urban planning, and drug discovery[27]. For example, a long-established antifungal drug, itraconazole, was found to act on the well-characterized Hedgehog pathway, particularly inhibiting the transmembrane receptor protein Smoothened. Independently, research has implicated Smoothened gain-of-function mutations as prevalent and key to the progression of certain cancer types such as lung, prostate, and skin. Once researchers connected these two disparate domains, trials were carried out and itraconazole is now an established chemotherapy potentiator[23].

Entity resolution, also known as record linkage or deduplication, is key to the success of link prediction due to the widespread integration of data from diverse sources. Entity resolution addresses the challenge of identifying and merging duplicate or related records between and within datasets, to enhance data quality, consistency, and accuracy. As datasets increasingly originate from heterogeneous sources, entity resolution becomes crucial for integrating disparate data and creating a unified, accurate representation of real-world entities[8].

We investigate recent advances in LLMs to improve link prediction via knowledge graphs generated from transdisciplinary literature. We investigate the challenges of zero-shot entity resolution using both ontology-based and large language model approaches. We study how workflow decisions affect the stability of link prediction results through a real-world application to research in chronic lower back pain.

## 2 Related Work

Within scientific natural language processing, resolving the ambiguity between important terms within both structured and unstructured forms of information is paramount for the collection of clean, consistent, and high-quality data. Methods of identifying related entities include Entity Linking, Entity Resolution, and Coreference Resolution.

Entity linking, also known as entity disambiguation, is the method of classifying the entities mentioned in unstructured text based on the terms inside a knowledge base [29]. Within entity linking, Broscheit (2014) showcases the potential of Pretrained Language Models by appending a classification layer on top of BERT in order to predict the probabilities of links existing between a given term and the entries within a knowledge base, i.e., Wikipedia [4]. Furthermore, Sakhovskiy et al. (2023) demonstrated the viability of knowledge graph-assisted entity linking with their Graph-Enriched Biomedical entity representation transformer, GEBERT [25].

Coreference Resolution is the method of identifying references within text that all allude to the same term [15]. In the field of Coreference Resolution, Li et al. (2021) injects domain-specific information from an external knowledge base into an LSTM [11] in order to improve Coreference Resolution on biomedical text [17]. Moreover, Lu et al. (2021) [18] compares the performance between methods specific to biomedical Coreference Resolution and language models pre-trained on biomedical text, such as BioBERT [16].

Entity Resolution, also known as Record Linkage, is the method of identifying mentions that refer to the same concept or entity throughout different sources and forms of data. [8]. In Entity Resolution, Huang et al. (2018) introduced a novel approach to the comparison and extraction of entities through the use of context graphs to evaluate and embed textual information to entities [14]. In addition, Obraczka et al. (2021) utilizes graph embedding similarity and graph attribute similarity as features for a machine learning classifier that predicts whether two entities match [22]. Although Entity Resolution, Entity Linking, and Coreference Resolution are all present in various portions of our LLM-driven text-to-knowledge graph workflow, this paper will focus mainly on entity resolution and how methods of merging identical mentions within our knowledge graph can impact methods such as link prediction.

Within Zero-shot Learning, Narayan et al. (2022) demonstrates the effectiveness of foundation models, such as GPT, on data preprocessing tasks such as Entity Matching, which seeks to identify identical records between datasets [20]. In biomedical applications of zero-shot learning, Hu et al. (2023) evaluates the potential of GPT models on clinical named entity recognition using zero-shot learning and prompt engineering [13]. In methods using ontologies, Prokofyev et al. (2015) uses ontologies to aid candidate clustering for coreference resolution. By assigning each mention an entity and type drawn from an ontology, the paper demonstrates how ontological annotations can improve clustering results [24]. In addition, Štajner et al. (2009) utilizes ontologies to generate a relatedness score to assist in entity resolution [26].

## 3 Methods

We focus the current work on the state of knowledge graph generation to discover and rank novel relationships between siloed areas of research. For our method, a research area is defined by its literature and a short description that defines the most relevant aspects of that literature. Therefore, our method takes as input two or more literature corpora with their associated descriptions and outputs a knowledge graph and predicted relations. While the way in which a corpus and description are defined is flexible, this paper formulates each description as a hypothesis-style paragraph that is then used to identify relevant literature. Once a description is written and a corpus of literature is defined, our knowledge graph construction and analysis workflow, outlined in Figure 1, can then be applied.

**Fig. 1:**
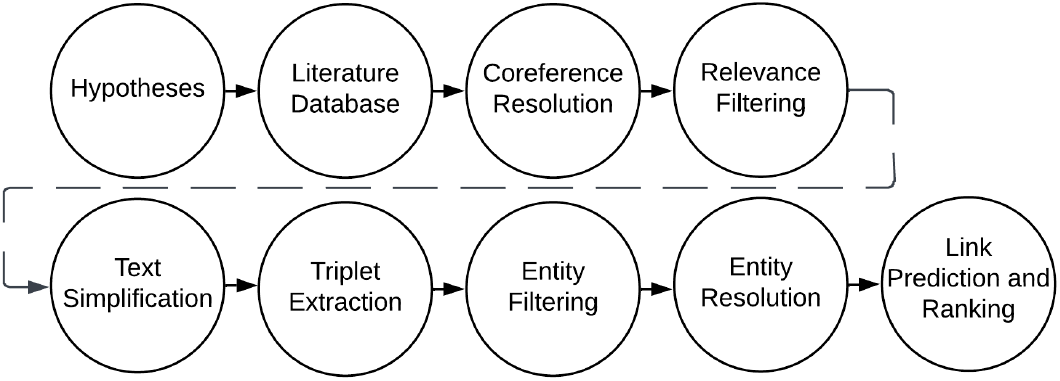
Overview of the link prediction and ranking workflow.

### 3.1 Hypotheses

We examined the performance of our workflow by measuring and discussing the utility of link prediction in the field of chronic lower back pain. We evaluated our workflow and its associated components on curated datasets created through the real-world application of relation discovery between two hypotheses derived from siloed areas of research in the field of chronic lower back pain. We formalize them as:

1. Cognitive therapy, mindfulness-based stress reduction, behavior therapy, and similar psychosocial-focused therapies produce similar pain reductions and improve other outcome measures. Hypothesis: most of the benefits derive from participating in an informative, interactive focused activity, rather than the distinguishing details of the specific treatment technique. [5]
2. Patients with cLBP exhibit altered gait patterns. The patterns are less complex compared with matched pain-free controls, presumably because pain anticipation and catastrophizing require high involvement of attentional processes during movement. Hypothesis: visually distracting cLBP patients will increase gait complexity and facilitate active rehabilitation. [12]

From each paper we procured PDFs of all references, barring paid textbooks and inaccessible journal articles, totaling 116 references across the two papers—40 of which pertain to the first paper [5], and 76 coming from the second paper [12]. The PDFs were compiled into a corpus for each respective hypothesis to be processed and extracted from. The first set of 40 references ranges across 24 different scientific journals. Most notably, 7 (17.5%) of them are from the Journal of Consulting and Clinical Psychology, another 7 (17.5%) are from The Journal of the International Association for the Study of Pain, and 3 (7.5%) are from The Clinical Journal of Pain. Meanwhile, the set of 76 references for hypothesis 2 spans 50 different journals, with 9 (11.8%) coming from the Journal of the International Association for the Study of Pain, 4 (5.3%) from Gait & Posture, and Chaos: An Interdisciplinary Journal of Nonlinear Science and the European Spine Journal contributing 3 (3.9%) articles each.

### 3.2 Workflow

In this section, we describe the details of our text-to-knowledge graph workflow, beginning after paragraphs are extracted from the articles described in Section 3.1.

#### Coreference resolution

First, our workflow preprocesses the text by utilizing SpanBERT [15], a language model that modifies BERT [10] by masking text segments, to perform coreference resolution. Afterward, the resulting text is passed toward relevance filtering.

#### Relevance filtering

In order to reduce computational workload and narrow our focus on the hypotheses, we utilize GPT for zero-shot relevance filtering. For each paragraph in the corpora, we query GPT to determine whether the text is relevant to either of the hypotheses using the prompt: “Your job is to read the following paragraph, and tell me whether it contains information relevant to the following hypothesis: [hypothesis text here]. Your response should be yes or no with no extra information.”.

#### Text simplification

The next step is to break each relevant paragraph into sentences which are then rewritten and simplified to aid triplet extraction. We query GPT to rewrite each sentence in the corpora into a simple subject-predicate form. This is done in two steps. First, we query GPT to label each sentence as simple, compound, complex, or compound-complex. Then, we make a second query to rewrite the non-simple text into simple sentences. These simplified sentences are then reformed into paragraphs and Coreference Resolution is performed again.

#### Triplet extraction

After simplifying the text, we extract the entities and relations using REBEL, a seq2seq language model that identifies both entities and relations concurrently [6]. An example sentence is shown in Figure 2. These triplets are then added to a knowledge graph and linked to their original sentences, paragraphs, and the hypothesis they refer to, capturing the provenance of the triplet. The head and tail of triplets are defined as instances of the *Entity* node class, while the relationship is defined as an edge between these two entity nodes.

**Fig. 2:**
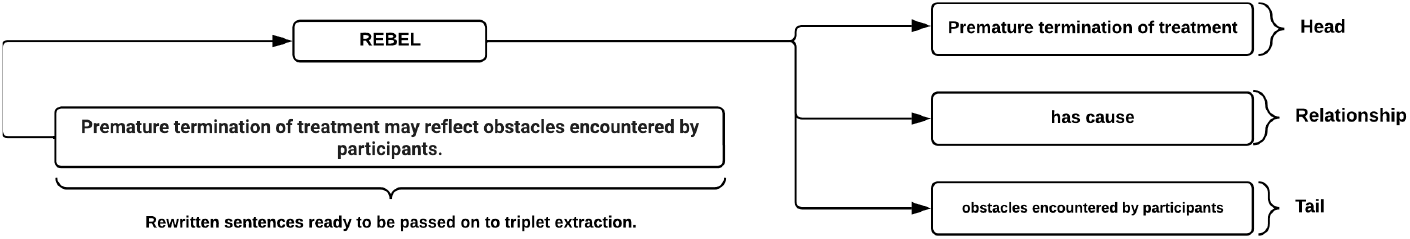
Example of triplet creation using REBEL on a GPT simplified sentence, extracted from Thorn et al. (2011) [28].

#### Entity Filtering

After extracting the entities and adding them to the knowledge graph, we filtered irrelevant terms to reduce the computational workload. In order to select terms that are relevant to the hypotheses, we started with the assumption that relevant entities could be recognized through their relationships defined by an ontology. To focus on triplets of relevance to the hypotheses, we manually extracted keywords from each hypothesis and annotated them using the NCIT ontology on the NCBO BioPortal. Triplets in the knowledge graph were likewise annotated by the same ontology, and triplets that shared an annotation with the keywords, were marked as relevant and used in future steps of the workflow.

#### Entity Resolution

The next step in the workflow is to identify pairs of entities equivalent in meaning. With this task arises two key challenges: The problem of evaluating ambiguous entities, that is, entities that have various meanings based on the context ascribed to them, and properly attributing the correct entities to their many entries across a corpus.

We implemented and evaluated three approaches on our labeled dataset. Those three approaches are based on the following technologies: (1) GPT, (2) Annotations via Ontology, and (3) GPT and Annotations via Ontology. The input to each algorithm is the head or tail phrase extracted from the REBEL model. These two inputs are then passed to the entity resolution algorithm, which returns True or False. If two nodes are determined to correspond to the same named entity, then the two nodes are linked with an edge. The first approach uses the GPT prompt: “You are a biomedical expert. Do the following mean the same thing? Your response should be yes or no with no extra information.”. The second approach looked at overlapping ontological annotations, and if the number of overlaps was greater than a threshold, two nodes were then linked. The third method was evaluated to improve computational efficiency, where the ontology annotation (second approach) is used to filter out unlikely matches, which are then passed to GPT (first approach).

#### Entity Merging

An entity resolution method resolves whether two entity nodes should be merged in a graph. This information is encoded by adding an edge between all pairs of equivalent nodes. Since more than two nodes may share the same meaning, there are multiple ways to determine which group of nodes should be merged together. If we assume that an edge indicates that two nodes are exactly equivalent in meaning, we can assume transitivity and consider any two nodes equivalent if there exists a path between them. Thus, we can merge all entities that belong to the same Weakly Connected Component (WCC) into a new node. However, in actuality, our automated methods may generate a loss of information when choosing to combine two entities. If we consider a graph (shown in Figure 3) such that edges *x* and *y* link nodes *a* and *b* as well as *b* and *c*, respectively, *a* and *b* may be equivalent and *b* and *c* may be equivalent, but *a* and *c* may not be equivalent. Thus, we consider two algorithms to help us identify these nodes: WCC and maximal cliques.

**Fig. 3:**
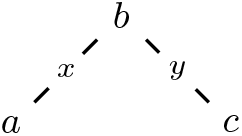
In this graph, WCC would identify*{a, b, c}* to be one component whereas maximal cliques would identify two components:*{a, b}* and*{b, c}*.

Nodes are in the same WCC if there exists a path between every pair of nodes (the direction of the edges is ignored). However, in the maximal clique algorithm, nodes are in the same clique if they share an edge with each other. For example, in Figure 3, WCC would identify nodes *a, b*, and *c* to all be in the same component. In contrast, maximal cliques would identify nodes *a* and *b* to be in one component and *b* and *c* to be a separate component.

After entity merging, we begin reconnecting the knowledge graph such that any edge *r* previously linking nodes *n* and *m*, where *n* is absorbed into *N* and *m* is absorbed into *M*, will now link nodes *N* and *M*. Thus, our relations remain consistent with our merged nodes.

#### Link prediction

We tested three link prediction algorithms. The first, *Nearest Neighbors* [21], captures that if vertex *x* and vertex *y* share many neighbors, then it is likely that they should be neighbors. A similar algorithm, preferential attachment, captures the idea that as networks grow, vertices that already have many neighbors are likely to gain more as time goes on [3]. Finally, we tested Adamic Adar which attempts to normalize for large disparities in edge counts between nodes [2].

### 3.3 Evaluation

#### Triplet Relevance Dataset

Using our workflow, we generated triplets from our corpora of papers related to hypothesis 1 and hypothesis 2. After grouping the triplets by their relation, we picked a single edge, displayed in Table 1, to analyze for downstream tasks such as link prediction. We observed that there were enough triplets connected by the “has cause” relation for data analysis, and we hypothesized that leveraging the “has cause” relation for link prediction would generate interesting discoveries. We extracted the triplets containing the “has cause” relation along with the sentence, the paragraph, and the hypothesis it originated from. After generating our dataset, we labeled each triplet as relevant to the source hypothesis, not relevant to the source hypothesis, or if an error was made in generating the triplet based on considering the head, tail, and whether the relation “has cause” was considered correctly applied.

**Table 1:** Counts of the top 20 most common relations extracted with REBEL.

| Edge Type | Count | Edge Type | Count |
| --- | --- | --- | --- |
| subclass of | 10016 | studied by | 680 |
| instance of | 5642 | use | 654 |
| facet of | 4749 | medical condition treated | 577 |
| part of | 3994 | followed by | 576 |
| has part | 3376 | studies | 565 |
| different from | 2024 | drug used for treatment | 551 |
| opposite of | 1613 | used by | 517 |
| has effect | 1334 | follows | 443 |
| has cause | 1005 | uses | 437 |
| said to be the same as | 974 | main subject | 418 |

#### Entity Resolution Dataset

After labeling the triplets, we extracted the samples marked as relevant and constructed a new set of entities defined as the union of all head and tail entities. Then, we generated all possible pairs of entities, filtered for duplicates, and stratified all pairs by their cosine similarity. Out of a dataset containing 1000 samples, we sampled 50 from a pool of samples with a cosine similarity of 1.0, 350 from a pool of samples with a cosine similarity less than 1.0 but greater than 0.9, and 600 from samples with a cosine similarity less than 0.9 and greater than 0.8. This new dataset was randomly condensed to 500 entity pairs, and labeled by multiple evaluators to introduce differing perspectives. Entity pairs were labeled as either matching, not matching, opposites, sub-classes, or linked. Examples of these types are shown in Figure 4. Afterward, we used the label data to benchmark our entity resolution methods.

**Fig. 4:**
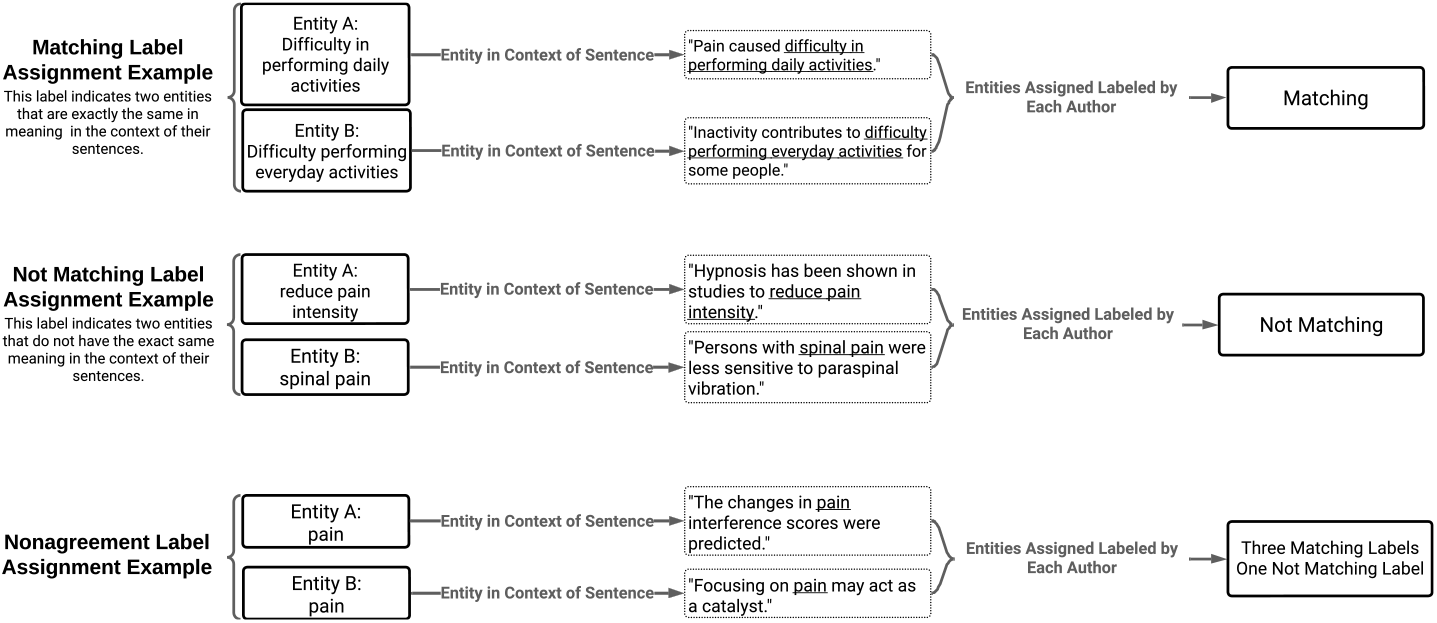
Entity Resolution Labeling Examples drawn from papers collected in our corpora.

## 4 Results and Discussion

### 4.1 Overview Statistics of Knowledge Graph

We first describe the overall statistics of the knowledge graph generated for our case study into transdisciplinary relation discovery for chronic lower back pain. After applying our workflow to the literature for each hypothesis, our knowledge graph contains the triple, paragraph, and sentence counts shown in Table 2. These results show that the number of extracted triplets increases with the number of sentences and paragraphs; however, we also see that there is significantly more data extracted for hypothesis 1 as opposed to hypothesis 2 across triplets, sentences, and paragraphs. As the size of the corpus for hypothesis 2 is larger than that of hypothesis 1, the number of paragraphs deemed relevant according to our filtering step is a smaller percentage of the total available. Finally, the keywords manually extracted from the hypotheses are shown in Table 3.

**Table 2:**
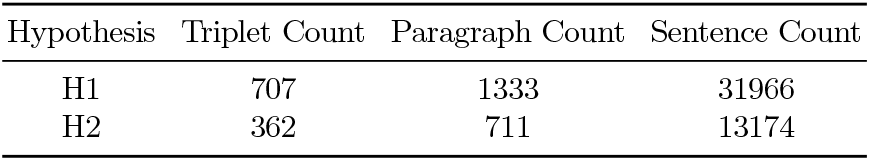
Triple, paragraph, and sentence count for each hypothesis based on the selected literature.

| Hypothesis | Triplet Count | Paragraph Count | Sentence Count |
| --- | --- | --- | --- |
| H1 | 707 | 1333 | 31966 |
| H2 | 362 | 711 | 13174 |

**Table 3:** Keywords manually extracted from each hypothesis to be used for entity filtering.

| Hypothesis 1 | Hypothesis 2 |
| --- | --- |
| treatment | active rehabilitation |
| interactive focused activity | gait complexity |
| informative activity | visual distractions |
| outcome measures | movement |
| pain reductions | attentional processes |
| similar psychosocial-focused therapies | catastrophizing |
| behavior therapy | pain anticipation |
| mindfulness-based stress reduction | pain-free controls |
| cognitive therapy | gait |
|  | gait patterns |

### 4.2 Labeled Datasets

We selected 1,053 triplets (head-relation-tail) for manual relevance labeling. Out of that total, 651 were labeled as relevant, 153 were labeled as erroneous (i.e., “has cause” not a correct relation), and 249 were labeled as not relevant. When labeling pairs of entities for equivalence, evaluators considered both the entity itself and the sentences where they were used. Evaluators, on average, labeled 13.25% of entity pairs as matching in meaning and context (Table 4). Strong agreement on matching entity pairs was rare: only 7.2% of entity pairs were labeled as matching by all 4 evaluators, and an additional 3.2% were labeled as matching by 3 out of the 4 evaluators (Table 5).

**Table 4:** Number of entity pairs labeled matching for resolution for each evaluator. Percents were calculated based on the total number of entities labeled per evaluator.

| Evaluator | Count labeled matching | Total labeled | Percent |
| --- | --- | --- | --- |
| 1 | 62 | 498 | 12.44 |
| 2 | 83 | 502 | 16.53 |
| 3 | 61 | 570 | 10.70 |
| 4 | 82 | 615 | 13.33 |
| Average |  |  | 13.25 |

**Table 5:**
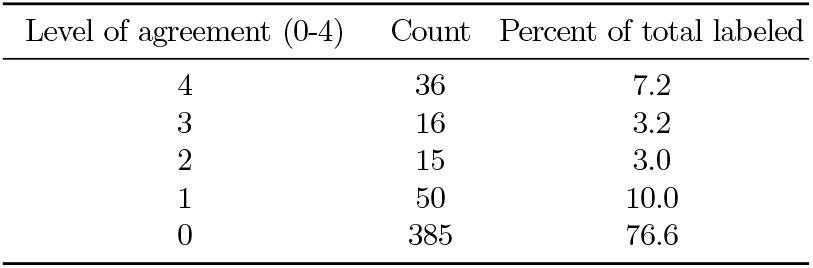
Number of evaluators that labeled a pair of entities as matching, where 4 out of 4 indicates strongest agreement. Percents were calculated as proportions of 500, the total number of entity pairs in this dataset.

### 4.3 Workflow Evaluation

The motivating task of this study is to identify relationships between and relevant to our two hypotheses. While manually labeling triplets and entities is ideal, this approach does not scale. Using the ontology as a bridge between our extracted triples and our hypotheses via entity filtering, we focused our analysis on 1,561 nodes that are involved in 292,269 relations.

We implemented and compared three entity resolution approaches. The results of these methods on our labeled dataset were used to determine the best method going forward. The outcomes of these three zero-shot algorithms are displayed in Table 6 and show that the Zero-shot GPT approach is effective in predicting which named entities should be combined when all four labelers are in agreement (Agreement = 1.0). The performance degrades as you consider pairs of nodes that received 75% agreement, and further declined when at or below 50% agreement. Furthermore, this GPT based approach took around half an hour to process approximately 500 pairs. In a deployed setting, entity resolution needs to consider many hundreds of thousands of pairs, making this approach a bottleneck in development and research.

**Table 6:**
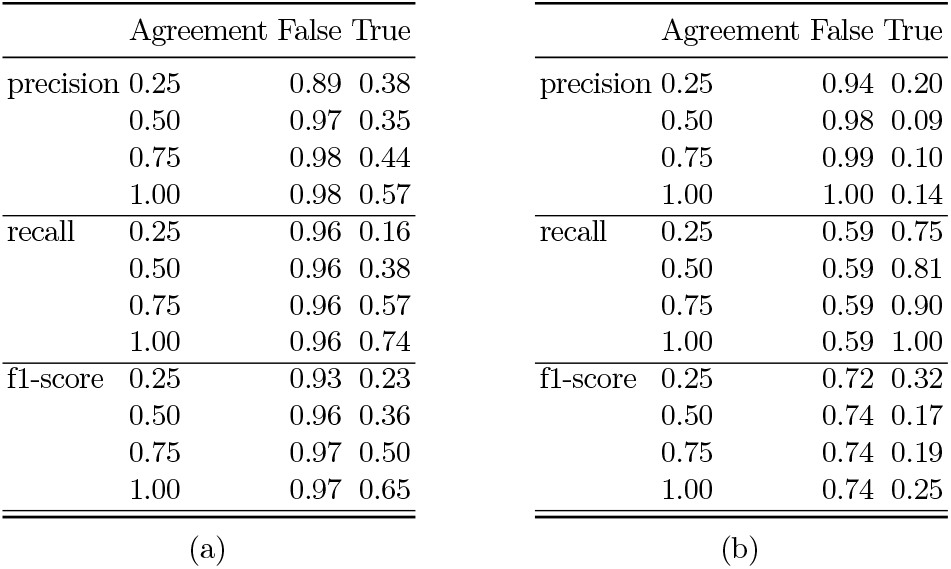
(a) GPT and (b) annotation entity resolution approach results grouped by agreement among labelers.

For approach 2 (ontology based), we explore annotations that overlap between terms using the BioPortal annotation API endpoint. Examining the number of overlapping annotations shared between pairs, we observed that nearly 300 of the 500 pairs share less than 5 annotations in common. For the purposes of evaluating this approach, we selected a threshold of 1 for the results shown in Table 6. These results indicate that an annotation approach can achieve high recall when attempting to identify potential nodes to combine; however, this is at the expense of precision. The running time of this approach is significantly shorter than the GPT approach and finished in less than 2 minutes. For this reason, we adopted the third approach, which was to first run the annotation-based filtering followed by the more precise but slower GPT approach.

The final step in the workflow is to merge nodes. For this task, we evaluated two strategies. The first was to merge all Weakly Connected Components, and the second was to merge only subsets of nodes that form a Maximum Clique. After entity resolution using approach 3 and applying WCC, our graph contains 1,229 merged nodes and 216 relations between them. Merging nodes using the maximum clique algorithm yields 1,423 merged nodes with 216 relations between them. Out of those 216 relations, both the WCC and maximum clique methods identified 39 “has cause” relations. To investigate the difference between the two methods, we compared merged nodes between the Weakly Connected Components and Maximum Clique approaches. Focusing our analysis on the nodes involved in a “has cause” relation, we found that 358 merged nodes were composed of the same entities in both the WCC and maximum clique methods. There were 33 merged nodes that differed between the two methods. For presentation purposes, we present a subset of 10 differing merged nodes in Table 7. Although our results indicate that applying WCC led to a more condensed graph as opposed to applying maximal cliques, our analysis indicates that the overall graph structure remains stable. Since applying Maximal Clique is far more computationally intensive than WCC, we decided to use WCC when constructing our knowledge graph for link prediction. The final step in our analysis is to compare and inspect the link prediction results.

**Table 7:** Differences between merged nodes using WCC and maximum cliques (subset of 10 shown)

| Text Description | WCC Node Count | Max Clique Count |
| --- | --- | --- |
| pain ratings | 0 | 4 |
| CBT intervention | 0 | 1 |
| Cognitive-behavioral therapy | 124 | 50 |
| pain intensity | 37 | 17 |
| pain-related fear | 0 | 4 |
| depression | 1 | 2 |
| psychological treatment | 7 | 15 |
| cognitively-focused CBT | 0 | 2 |
| cognitive behavioral therapy | 12 | 18 |
| Depression | 3 | 2 |

For this, we ran the Adamic Adar, preferential attachment, and nearest neighbors link prediction algorithm after merging the nodes using the WCC approach. The results between preferential attachment and nearest neighbors are nearly identical, with a Kendall rank correlation coefficient of 0.94. Adamic Adar produced significantly different results than both preferential attachment and nearest neighbors with correlation coefficients of -0.05 and -0.16, respectively. Due to Adamic Adar normalizing for edge disparities, links predicted by Adamic Adar may reflect more nuanced relations between entities. However, links predicted by preferential attachment and nearest neighbors are likely going to be easier to verify in existing literature that may currently not be reflected in the graph. Thus, we chose to analyze the results of nearest neighbors, where Table 8 shows the top 20 predicted relations. Nearest neighbors produced 1,128 relations with a score of *>*0 out of a possible 2,231 pairs of nodes.

**Table 8:** Using the WCC merged nodes, link prediction results nearest neighbors.

| Node A Description | Node B Description | Score |
| --- | --- | --- |
| Cognitive-behavioral therapy | pain | 215 |
| pain | pain intensity | 165 |
| Cognitive-behavioral therapy | pain intensity | 154 |
| pain | Chronic pain | 142 |
| pain | pain interference | 140 |
| Cognitive-behavioral therapy | Chronic pain | 131 |
| Cognitive-behavioral therapy | pain interference | 128 |
| pain | Pain | 127 |
| pain | pain catastrophizing | 126 |
| pain | pain-related distress | 124 |
| pain | treatment | 124 |
| pain | Behavioral therapy | 119 |
| pain | decrease in depressive symptoms | 117 |
| pain | positive changes | 116 |
| pain | cognitive process | 116 |
| pain | BIO treatment condition | 116 |
| pain | pain-related behaviors | 116 |
| pain | pain symptoms | 116 |
| pain | body image disruption | 116 |
| Cognitive-behavioral therapy | Pain | 116 |

Some notable predicted links emerge in the top 20, such as “pain” and “decreases in depressive symptoms” or “pain” and “body image disruptions”. While the significance of these associations is unknown, we are impressed by the ability of the workflow to span disparate fields (ex: pain as a clinical feature and insights into psychological features). Looking back at the hypotheses and their associated references, we can see where this link could have been initiated, especially in consideration of entity resolution and ontology processing. However, we argue that without the workflow, this novel connection could have been missed therefore limiting potentially important chronic pain diagnostics or treatments. We do not propose that these associations do not warrant future scrutiny or are without critique. We are instead intrigued by the possibility of novel connections between previously siloed fields being brought together out of the noise of their respective fields to be investigated by domain experts

## 5 Future work

Throughout the development of our workflow, we made numerous decisions to optimize running time and to focus our analysis on entity resolution. This leaves many avenues of refinement open for future work. Our workflow can be upgraded as large language models improve. Furthermore, we focused our evaluation on entity resolution and link prediction, and similar treatment for sentence simplification, filtering, etc would improve the workflow. Specifically to entity resolution, it will be interesting to explore methods of incorporating the sentence of each entity to match context-dependent pairs as this was important to our manual labelers. After experimenting with entity resolution, further analysis of the performance differences between Weakly Connected Components and Maximal Cliques will be interesting to explore. In addition, we are interested in measuring the impact of reframing our entity resolution problem as whether clusters or subgraphs of entities refer to the same term instead of pairs of entities. Going forward, our workflow will serve as a foundation for studying methods of knowledge graph construction that best enable link prediction. We would like to further test the stability and reproducibility of this pipeline, especially in the context of the sparseness and incomplete data prevalent in electronic health records. We hypothesize that persistent link predictions emerging despite sparse data will be an additional metric to support the validity of the novel associations. We believe this iteration of the workflow is an important first step in capitalizing on the power of knowledge graphs and large language models in generating important new connections between complicated and siloed biomedical fields.

## 6 Conclusion

In this study, we presented an end-to-end knowledge graph construction workflow and demonstrated its capabilities in predicting links between two different hypotheses within chronic pain. We displayed each step of our workflow, including Coreference Resolution, Relevance Filtering, Text Simplification, Triplet Extraction, Triplet Filtering, Entity Resolution, and Entity Merging, and analyzed how the decisions made in the workflow impacted our link prediction results. In addition, we also presented the Triplet Relevance dataset, a benchmark to evaluate methods of filtering irrelevant information, and the Entity Resolution dataset, a benchmark to test methods of identifying entities of equivalent meaning. In the curation of these datasets, we recognized that entity resolution is an extremely difficult task even for domain experts, as illustrated by the seldom agreement of all evaluators. In an attempt to model the evaluators, we observed that GPT outperforms the use of an ontology, although far less efficiently in terms of computation. Afterward, we applied two separate approaches: Weakly Connected Components and Maximal Clique. Since we observed negligible differences between Weakly Connected Components and Maximal Clique, we used WCC to merge nodes and finalize our knowledge graph ahead of performing link prediction. To that extent, various methods of link prediction were utilized which gave insight as to why a domain expert may select one method over another. Our workflow found potential connections between disparate fields. We believe this type of automated link prediction can find critical connections between complicated fields, potentially improving clinical diagnostics and care. Our code and workflow is available at https://github.com/calpoly-bioinf/iwbbio_2024.

## 7 Acknowledgments

We thank members of the Cal Poly Bioinformatics Research Group for their invaluable help with this research. This work was supported by “UCSF Core Center for Patient-centric Mechanistic Phenotyping in Chronic Low Back Pain (UCSF REACH)” funded by NIH. Support was provided for the Computational Molecular Sciences Center by the Frost Fund at the Cal Poly Bailey College of Science and Math.

